# VIGA: an one-stop tool for eukaryotic Virus Identification and Genome Assembly from next-generation-sequencing data

**DOI:** 10.1101/2023.06.14.545025

**Authors:** Ping Fu, Yifan Wu, Zhiyuan Zhang, Ye Qiu, Yirong Wang, Yousong Peng

**Affiliations:** Bioinformatics Center, College of Biology, Hunan Provincial Key Laboratory of Medical Virology, Hunan University, Changsha 410082, China

**Author notes:** To whom correspondence should be addressed. (YP).

## Abstract

Identification of viruses and further assembly of viral genomes from the next-generation-sequencing (NGS) data are essential steps in virome studies. This study presented an one-stop tool named VIGA (available at https://github.com/viralInformatics/VIGA) for eukaryotic virus identification and genome assembly from NGS data. It was composed of four modules including identification, taxonomic annotation, assembly and novel virus discovery which integrated the homology-based method for virus identification and both the reference-based and *de novo* assemblers for accurate and effective assembly of virus genomes. Evaluation on multiple simulated and real virome datasets showed that VIGA assembled more complete virus genomes than its competitors on both the metatranscriptomic and metagenomic data, and also performed well in assembling virus genomes at the strain level. Finally, VIGA was used to investigate the virome in metatranscriptomic data from the Human Microbiome Project and revealed different composition and positive rate of viromes in diseases of Prediabetes, Crohn’s disease and Ulcerative colitis. Overall, VIGA would help much in identification and characterization of viromes in future studies.

## Introduction

Viruses are ubiquitous in nature and have profound impacts on the health and diversity of all living organisms^[1]^. However, most viruses remain unknown and unculturable due to the limitations of conventional isolation methods^[2]^. In recent years, the rapid development of next-generation-sequencing (NGS) technologies has revolutionized the field of metagenomics and transcriptomics^[3]^, enabling the analysis of nucleic acid sequences obtained directly from environmental or clinical samples. For example, the Human Microbiome Project^[4]^ has facilitated the generation of hundreds of reference microbial genomes, along with thousands of metagenomic and metatranscriptomic datasets from multiple parts of the human body. This has greatly increased the number of viral genomes and provided valuable information for studying viral evolution, diversity and epidemiology^[5]^.

Identification of viruses from NGS data is the first step in virome studies. There are currently two kinds of methods for virus identification. The first is the homology-based methods, such as using tools of BLAST^[6]^ or HMMER^[7]^ to search for viruses which have sequence similarity to known viruses. This kind of methods is commonly used in identifying eukaryotic viruses which have been studied much. For example, Moustafa et al.^[8]^ identified 19 viruses by whole genome sequencing of blood from 8240 individuals using BLAST. The advantage of these methods is that they are generally accurate with a low rate of false positives^[9]^, but they may miss identification of novel viruses which have remote or little homology with known viruses^[10]^. This problem becomes serious in discovery of phages which have huge genetic diversity. To address the problem, the other kind of methods has been developed for virus identification based on machine-learning methods, such as Virtifier^[11]^, Seeker^[12]^ or VirFinder^[13]^. They can detect viruses with higher sensitivity than the homology-based methods and are generally used in identifying phages. For example, the project of Global Ocean Viromes 2.0 (GOV 2.0)^[14]^ have identified 195,728 viral populations from the global ocean DNA virome dataset based on this kind of methods. Unfortunately, a high false positive rate was observed for these methods^[15]^ and there was also inconsistency between the prediction results by different methods^[16]^.

Assembling virus genomes is also an essential step for further studies of the virome. Unfortunately, assembling large genomic fragments from short-read sequencing data is a formidable computational challenge^[17]^. In the case of viruses, the presence of repetitive region^[18]^, strain heterogeneity^[19]^ and low-abundance viral populations^[20]^ can pose significant difficulties for accurate assembly of virus genomes. Besides, the quality of the assembly may be compromised by errors in the sequencing data. To address these challenges, researchers have developed various methods for improving the accuracy and quality of the virus genome assembly which can be mainly grouped into two categories^[21]^: one is the reference-based assembly methods, such as MetaCompass^[22]^ and VirGenA^[23]^ and the other is the *de novo* assembly methods, such as Trinity^[24]^ and Haploflow^[25]^. The reference-based methods assembling genomes by taking a known genome as a guide. The advantage of this kind of methods is that they are generally more accurate than the *de novo* methods^[22]^, but they are not suitable for genome assembly of viruses without reference genomes. On the contrary, the *de novo* methods can be applied to all viruses including novel viruses, although they generally require a deep sequencing depth and may not assemble complete genomes^[26]^.

In this study, an one-stop tool named VIGA was developed for eukaryotic virus identification and genome assembly from NGS data. It combines both functions of virus identification and genome assembly. Multiple assembly tools including both the reference-based and *de novo* ones were integrated in VIGA for accurate and effective assembly of virus genomes. Evaluation on multiple simulated and real virome datasets showed that VIGA could be used for assembling virus genomes and separating mixtures of virus strains from metagenomic and transcriptomic data.

## Materials and methods

### Data collection

To evaluate the performance of computational tools in virus identification and genome assembly, three datasets were used : the first is a dataset of mock viral community (Accession in NCBI BioProject database^[27]^: PRJNA431646^[17]^) which consisted of 3,090,013 paired-end sequencing reads from seven viruses with varying proportions: human poliovirus 1 (47.04%), human mastadenovirus C (30.90%), coxsackievirus B4 (13.30%), murid gammaherpesvirus 4 (7.43%), echovirus E13 (0.78%), human mastadenovirus B (0.55%) and rotavirus A (0.001%).

The second is an RNA-Seq dataset of sweet potato viromes which included 10 samples (Accession in NCBI BioProject database: PRJNA517178^[28]^). Previous studies have identified ten virus species including the sweet potato symptomless virus 1 (SPSMV), sweet potato latent virus (SPLV), sweet potato virus G (SPVG), sweet potato virus 2 (SPV2), sweet potato virus C(SPVC), sweet potato leaf curl virus (SPLCV), sweet potato feathery mottle virus (SPFMV), sweet potato virus E(SPVE), sweet potato virus F (SPVF), sweet potato chlorotic fleck virus (SPCFV) from the dataset using the homology-based method, and also obtained their genomes (Accession numbers in NCBI GenBank database: NC_034630, NC_020896, NC_018093, NC_017970, NC_014742, NC_004650, NC_001841, NC_006550, MH388501 and MH388502) by the RT-PCR method.

The third is a viral metagenomic dataset of the bird fece which included 14 samples (Accession in NCBI SRA database: SRR10873474, SRR10874379, SRR10874826, SRR10873746, SRR10873876, SRR10873966, SRR10874819, SRR10874825, SRR10875205, SRR10873770, SRR10873785, SRR10873756, SRR10874841, SRR10874829^[29]^). Previous studies have identified 16 viral strains of three virus species including Riboviria sp. (Accessions of viral genome sequences in NCBI GenBank database: MN933892.1, MT138199.1, MT138205.1, MT138191.1, MN933887.1, MT138420.1 and MT138407.1), CRESS virus sp. (Accessions: MN928923.1, MN928933.1, MN928938.1, MN928948.1, MN928929.1, MT138070.1 and MT138040.1) and Picornavirales sp. (Accessions: MT138390.1 and MT138137.1) from the dataset using the homology-based method, and also obtained virus genomes by the RT-PCR method.

To evaluate the performance of computational tools in assembling viral genomes at the strain level, two datasets were used. The first is the HIV dataset which was created in Fritz’s study by mixing three HIV strains with about 95% sequence identity in proportions 10:5:2^[25]^. The dataset contains simulated metagenomic sequencing reads with the length of 150 bp and the depth of 20,000 (available for download at https://frl.publisso.de/data/frl:6424451/). The other is the HBV dataset^[30]^ which is the metagenomic sequencing of two clinical samples that were co-infected by two HBV strains with 89% sequence identity (Accessions in NCBI SRA database: ERR3253398 and ERR3253399).

The metatranscriptomic datasets derived from the Human Microbiome Project (https://portal.hmpdacc.org/)^[4]^ were used to illustrate the usage of VIGA. Only the paired-end sequencing samples were used as VIGA can only use the paired-end data. A total of 1321 samples were finally used. The meta information including the disease state, age and gender of samples was also obtained (Table S1).

### The workflow of VIGA

VIGA included four modules: Identification, Taxonomic annotation, Assembly and Novel Virus Discovery which were described as follows. The Identification module aimed to detect eukaryotic virus sequences from NGS data. Firstly, fastp (version 0.21.0)^[31]^ was used to trim the universal adapter sequences from raw reads and filter reads with average base quality less than Q20. Secondly, the remaining clean reads were assembled into contigs using the Trinity program (version 2.1.1)^[24]^ with default parameter settings. Thirdly, the contigs were queried against a library of virus protein sequences retrieved from the NCBI RefSeq database on June 10^th^, 2020 using Diamond BLASTX (version 0.9.25.126)^[32]^. The contigs with an E-value of less than 1E-5 to the best hit were labeled as hypothetical viral contigs. Fourthly, the hypothetical viral contigs were queried against the NR database (downloaded on November 17^th^, 2020) with Diamond BLASTX to remove false positives. Those with the BLASTX best hit belonging to viruses were kept and considered as viral contigs. Finally, they were filtered based on host of the BLASTX best hit, and only those with the BLASTX best hit belonging to eukaryotic viruses were kept for further analysis.

The Taxonomic annotation module was designed for taxonomic annotation of viral contigs identified above. The taxonomy information of viral contigs was obtained according to the identity and coverage between the contig and the BLASTX best hit against the NR database. If the identity and coverage were no less than 90% and 80%, respectively, the viral contig was considered to be the same virus species with the BLASTX best hit; if they were no less than 70% and 60%, respectively, the viral contig was considered to be the same genus with the BLASTX best hit; if they were no less than 50% and 40%, respectively, the viral contig was considered to be the same family with the BLASTX best hit. For viral contigs which were annotated at the species level, the Assembly module would be conducted; for those which were annotated at the genus level, the Novel Virus Discovery module would be conducted.

The Assembly module assembled and quantified the viral genome for the virus species after the taxonomy annotation. Firstly, a library of virus reference genomes was built as follows: i) genome sequences of all eukaryotic viruses were downloaded from the NCBI Genome database on August 17^th^, 2022. ii) genome sequences of the same virus species were clustered using the MMseqs2 (version 13.45111)^[33]^ at 100% level. The representative sequence in each cluster was added to the library of virus reference genomes. Secondly, all representative sequences of the virus species were taken as reference genomes and were used to assemble the virus genome using MetaCompass (version 2.0.0, parameter: ‘-l 100 -t 30’)^[22]^. Thirdly, RagTag (version 2.1.0, parameter: ‘-u -C’)^[34]^ was used to correct potential assembly errors in contigs based on the reference genomes. Then, MetaQUAST (version 5.0.2)^[35]^ was used to evaluate the quality of assembled viral genomes and output the Genome Fraction of the assembled virus genome which measures the genome completeness. Then, the assembled viral genomes were quantified using the FPKM (fragments per virtual kilobase per million sequenced reads) method according to Lee’s study^[36]^ which was listed as follows:

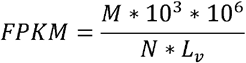

where L_v_ is the length of the assembled viral genome (bp), M is the number of reads assigned to the virus genome, N is the total number of clean reads. The Genome Coverage^[37]^ and the Depth Coverage^[38]^ of the assembled genome which measure the sequencing broadness and depth of the genome, respectively, would also be outputted. Finally, the Depth Coverage of the assembled genome would be visualized with a line chart.

The Novel Virus Discovery module was designed for novel virus discovery. Viral contigs which were annotated as the same genus were pooled together and defined as a new virus class (NVC). The Read Coverage of the NVC which measures the sequencing depth of the NVC, the identity and coverage between viral contigs and the BLASTX best hits against the NR database, and the detailed information of the BLASTX best hits would be provided.

### Comparison of VIGA and other assembly tools

Four commonly used assembly tools including two reference-based ones, i.e., MetaCompass^[22]^ and VirGenA^[23]^, and two *de novo* ones, i.e., Haploflow^[25]^ and Trinity^[24]^, were used for virus genome assembly and for comparison to VIGA. MetaCompass has been reported to have high accuracy in resolving complex and highly variable regions in genome assembly. The local version of MetaCompass (version 2.0.0, available at https://github.com/marbl/MetaCompass) was used by specifying the reference genome and setting the read length of 100 and other parameters as default. VirGenA has been optimized for viral genome assembly from low-coverage datasets. The local version of VirGenA (version 1.4, available at https://github.com/gFedonin/VirGenA/releases) was used with the insertion length of 800 and other parameters as default. Haploflow is a state-of-the-art *de-novo* assembly tool that can accurately reconstruct haplotype genomes from sequencing data. The local version of Haploflow (version 1.0, available at https://github.com/hzi-bifo/Haploflow) was used with default parameters; Trinity is a de Bruijn graph-based method for efficient and robust *de novo* reconstruction of transcriptomes from RNA-Seq data. The local version of Trinity (version 2.1.1, available at https://github.com/trinityrnaseq/trinityrnaseq) was used with default parameters.

The Genome Fraction (GF)^[35]^ was used to measure the completeness of the assembled genome. It was calculated as the ratio of the virus genome covered by the bases to which at least one contig has alignment. Two additional metrics were used to measure the accuracy and precision in assembling virus genomes at the strain level. One is the Mismatches per 100 kb (M100K)^[35]^ which referred to the average number of mismatches per 100,000 aligned bases and was used to measure the accuracy of assembled virus genomes. The other is the Strain Precision (SP)^[25]^ which was calculated as the ratio of correctly assembled contigs among all contigs of the virus and was used to assess the precision of assembled virus genomes.

### Virus identification and genome assembly on the Human Microbiome Project

VIGA was used to identify viruses and assemble virus genomes from the Human Microbiome Project. The host kingdom (plant, fungi and animal) of the identified viruses was inferred based on the virus family as viruses of the same family generally infect host of the same kingdom. When analyzing the virome in diseases, only animal viruses with abundance of greater than 2 FPKM were kept. Besides, viruses of the Retroviridae family were removed due to possible contamination from endogenous retroviral sequences, and viruses of the Baculoviridae family were also removed which are commonly used in the laboratory.

### Statistical analysis

The Pearson Correlation Coefficient (PCC) was used to measure the correlation between virus abundance and virus percentage in the mock virus community. Python (version 3.8) was used for statistical analysis.

## Results

### Overview of the VIGA pipeline

The VIGA included four modules: Identification, Taxonomic annotation, Assembly and Novel Virus Discovery (Figure 1) which were described briefly as follows. The Identification module aimed to detect eukaryotic virus sequences from NGS data. Firstly, the raw reads were pre-processed and then were assembled into contigs. Then, these contigs were queried against virus protein sequences using BLAS<u>TX</u>, and candidate virus contigs were obtained. Subsequently, to remove false positives, the candidate virus contigs were further queried against the NR database using BLASTX. Finally, the virus contigs were filtered based on host, and only eukaryotic virus sequences were kept for further analysis.

**Figure 1.**
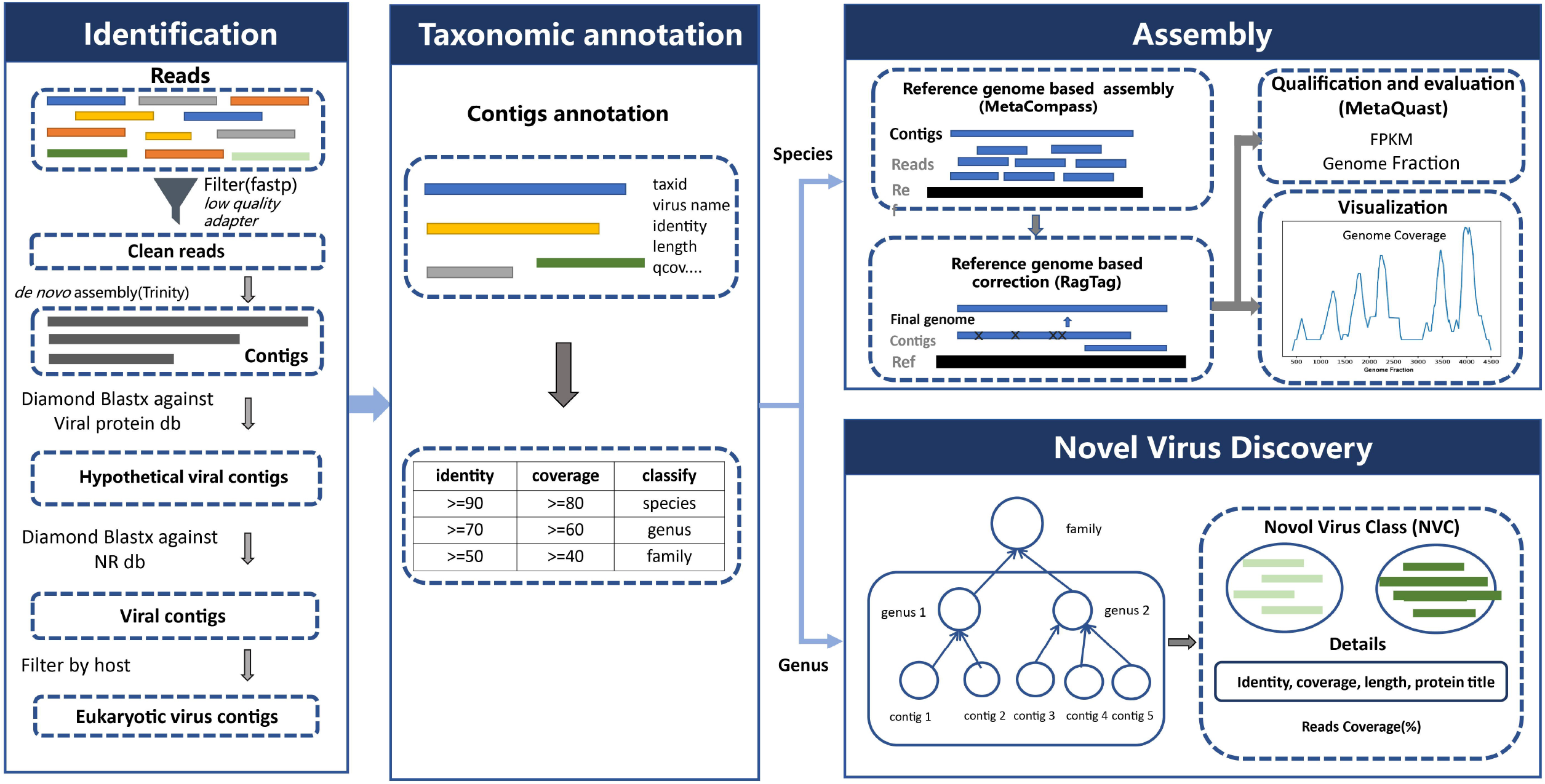
The workflow of VIGA.

The Taxonomic annotation module was designed for taxonomic annotation of viral contigs identified above. The taxonomy information of viral contigs was obtained according to the identity and coverage between the contig and the BLASTX best hit against the NR database (see Materials and methods). For viral contigs which were annotated at the species level, the Assembly module would be conducted; for those which were annotated at the genus level, the Novel Virus Discovery module would be conducted.

The Assembly module assembled and quantified the viral genome for the virus species after the taxonomic annotation. Firstly, the reference genomes were obtained for the virus species; then, MetaCompass was used to assemble the virus genome based on reference genomes of the virus species; then, RagTag was used to assembly the virus genome after correcting potential assembly errors in contigs based on the reference genomes; finally, the assembled viral genomes were evaluated with MetaQUAST and quantified using FPKM.

The Novel Virus Discovery module was designed for novel virus discovery. Viral contigs which were annotated as the same genus were pooled together and defined as a new virus class (NVC). Then, the Read Coverage of the NVC, the identity and coverage between viral contigs and BLASTX best hit against the NR database, and the detailed information of the BLASTX best hits were provided.

### Evaluating the performance of VIGA on a mock viral community

The performance of VIGA was firstly evaluated on a mock viral community which consisted of seven viruses with different proportions (Materials and methods). For comparison, we also evaluated four commonly used tools including two reference-based (MetaCompass and VirGenA) and two *de novo* tools (Trinity and Haploflow). As illustrated in Figure 2A, VIGA successfully identified six viruses except the rotavirus A. For viruses of PV-1, EV-13, HAdVC, and CV-B4, the Genome Fractions of viral genomes assembled by VIGA exceeded 0.94 which were much higher than those of its competitors. For example, the Genome Fractions of viral genomes assembled by MetaCompass and Trinity ranged from 0.74 to 0.94, 0.50 to 0.84, respectively. For viruses of HAdV-5 and HAdV-11, VIGA performed poorly, with the Genome Fractions of 0.636 and 0.064, respectively; however, it still outperformed its competitors much.

**Figure 2.**
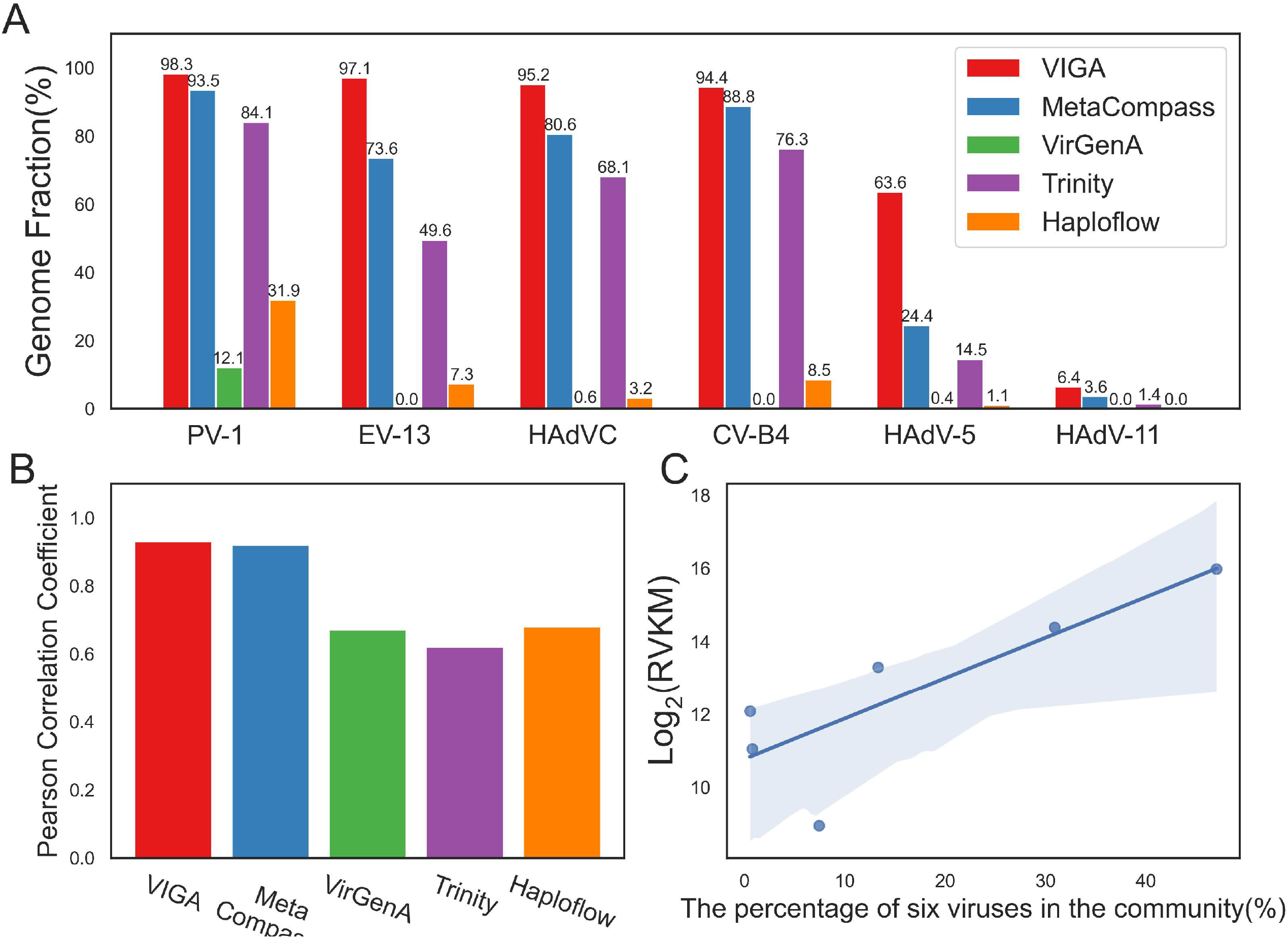
The performances of VIGA and its competitors on a mock virus community. (A) The ratios of viral genomes assembled by VIGA and its competitors including MetaCompass, VirGena, Trinity and Haploflow. (B) The Pearson Correlation Coefficient between the calculated virus abundance based on the viral genomes assembled by different software tools and the viral percentages in the virus community which reflect the real virus abundance. (C) The scatter plot of the calculated virus abundance based on VIGA and the real virus abundance. The blue line referred to the linear regression and the grey area referred to 95% confidence interval. PV-1, the human poliovirus 1; EV-13, the human echovirus 13; HAdVC, the human adenovirus C; CV-B4, the coxsackievirus B4; HAdV-5, the human adenovirus 5; HAdV-11, the human adenovirus type 11.

Previous studies have shown that the transcript abundance which was calculated based on the effective length (defined as the length of transcript regions that were covered by uniquely mapped reads) can reflect the gene expression more accurately than the abundance based on the full length of the transcript. Hence, the length of the assembled virus genome was taken as the effective length and was used to calculate the virus abundance using the FPKM method (Materials and methods). The Pearson Correlation Coefficient (PCC) was used to measure the correlation between the calculated virus abundances and the percentages of viruses in the mock virus community which reflect the real virus abundance. As shown in Figure 2B&C, VIGA had a PCC of 0.93 which was similar to that of MetaCompass (0.92) and much higher than those of other software tools (0.62-0.68), suggesting that the virus quantification based on the virus genome assembled by VIGA could accurately capture the real virus abundance in complex virus communities.

### Performance of VIGA on metatranscriptomic and metagenomic datasets

Then, VIGA was evaluated on a metatranscriptomic dataset of the sweet potato virome from which previous studies have identified ten viruses and obtained their genomes by the RT-PCR method. VIGA and other four assemblers (MetaCompass, VirGena, Trinity and Haploflow) were used to assemble virus genomes in the dataset. Except for SPCFV, all viruses were assembled by at least one tool. The Genome Fraction of virus genomes assembled by VIGA ranged from 1.6% to 100% for these nine viruses (Figure 3A), with a median of 47.9% which was greater than those of other software tools (median ratio ranging from 0% to 27.1%). Notably, both VIGA and MetaCompass assembled near complete genomes for five viruses including sweet potato virus G (SPVG, 99.17%), sweet potato leaf curl virus (SPLCV, 100%), sweet potato feathery mottle virus (SPFMV, 99.89%), sweet potato virus E (SPVE, 99.94%) and sweet potato virus F (SPVF, 99.45%), three of which (SPLCV, SPFMV and SPVE) were also assembled well by Trinity. However, both VirGena and Haploflow performed poorly and assembled only a small proportion of genomes for all nine viruses.

**Figure 3.**
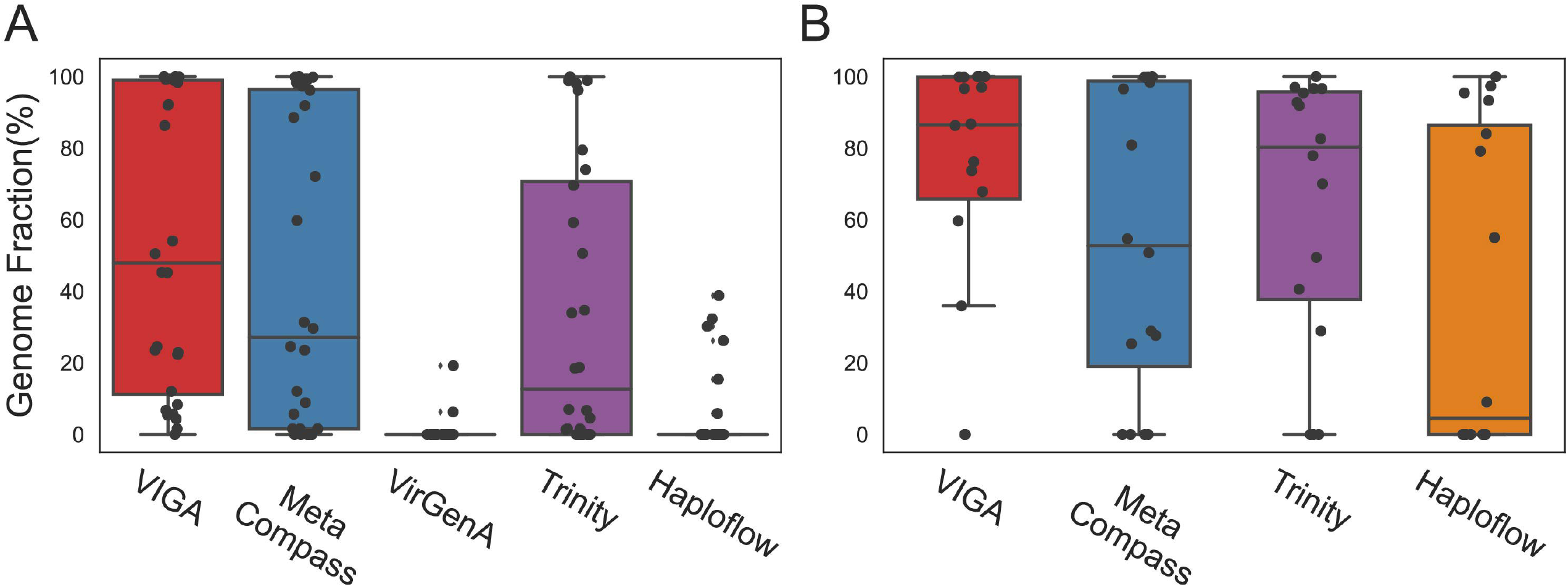
Performance comparison of different software tools on metatranscriptomic (A) and metagenomic (B) datasets. Boxplots show the statistical distribution of Genome Fraction of assembled virus genomes by different tools. Each dot in the boxplots referred to the Genome Fraction of the virus genome assembled in a sample. The results of VirGenA on the metagenomic data were not shown as no viral genomes were assembled sucessfully by the assembler.

VIGA was also evaluated on a metagenomic dataset of bird droppings from which 16 viral strains of three virus species were identified and their genomes were obtained by the RT-PCR method. VIGA stood out as the most effective tool for assembling viral genomes, with the Genome Fraction ranging from 0% to 100% and a median of 86.54% (Figure 3B), while other three assemblers (MetaCompass, Haploflow and Trinity) had the median Genome Fractions ranging from 4.5% to 80.3%. Notably, VIGA assembled high ratios of virus genomes in most samples, with only two exceptions resulting in no genome assembly.

### Performance of VIGA in assembling virus genomes at the strain level

We then evaluated the performance of VIGA and its competitors in assembling virus genomes at the strain level. Two datasets were used in the evaluation. The first is the HIV dataset which was the simulated metagenomic sequencing of three HIV strains with a 95% sequence identity. VIGA assembled genomes of all three strains successfully. Three indexes including Genome Fraction (GF), Strain Precision (SP) and Mismatches per 100 kbp (M100K) were used to measure the performance of assemblers (see Materials and methods). As shown in Figure 4A and Table S2, VIGA achieved near perfect performance on indexes of GF and SP, with an average GF of 98.2% and a SP of 100%, while in terms of M100K, VIGA had 2787 mismatches per 100 kbp which was much smaller than those of other assemblers except Haploflow that had the lowest mismatches per 100 kbp (33.7) among all assemblers. Compared to VIGA, Haploflow also achieved excellent performances on these indexes, with the SP of 100% and the average GF of 93.4%; other three assemblers had medium or poor performances on at least one index. For example, VirGenA had poor performances on the GF (76.1%).

**Figure 4.**
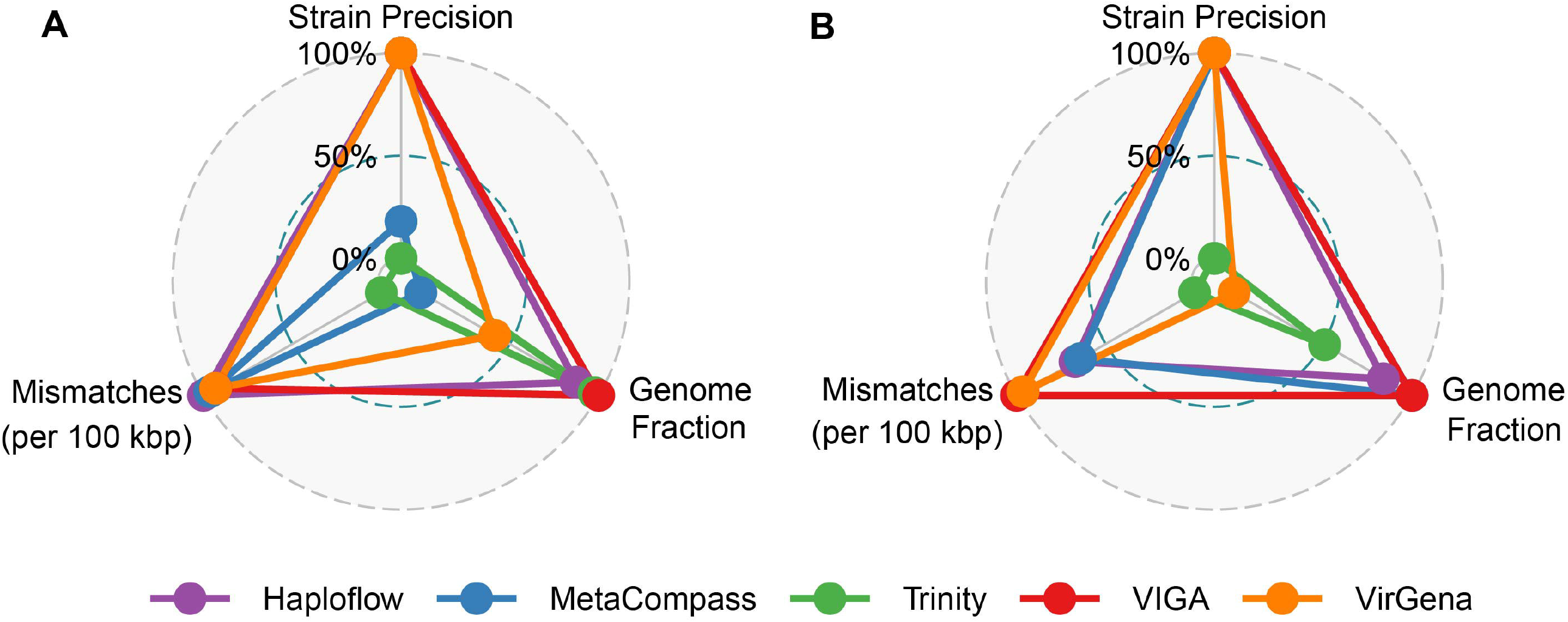
Performance of five assemblers (Haploflow, MetaCompass, Trinity, VIGA and VirGena) on the HIV (A) and HBV (B) datasets. Three indexes including Genome Fraction (GF), Strain Precision (SP) and Mismatches per 100 kbp (M100K) were used to measure the performances of assemblers. Each index was normalized with the Min-Max Normalization method before it was used in the radar plot.

The second is the HBV dataset which was the metagenomic sequencing data of two samples infected by two HBV strains with 89% sequence identity. As shown in Figure 4B and Table S3, VIGA performed best on three indexes among all assemblers, with the average GF of 99.9%, SP of 100% and 1890.7 mismatches per 100 kbp; Haploflow performed secondly among all assemblers, with the average GF of 91.2%, SP of 100% and 2355 mismatches per 100 kbp; other assemblers had median or poor performances on at least one index. For example, Trinity had an average GF of 73.5% and 3306.4 mismatches per 100 kbp.

### Application of VIGA in identifying and assembling virus genomes from the Human Microbiome Project (HMP)

In order to illustrate the practical usage of VIGA on large datasets, we reanalyzed 1321 metatranscriptomic samples from the HMP project (https://portal.hmpdacc.org/) using VIGA, with the total data volume of 1.14T. A total of 125 known eukaryotic viruses were identified from 467 samples, with 72 plant viruses, 31 animal viruses, 4 fungal viruses and 18 viruses infecting both plants and fungi (Figure 5A) (Table S4). The Genome Fractions of the assembled virus genomes ranged from 1.4% to 100% with a median of 33.8% (Figure 5B). A total of 44 viruses were assembled with high completeness (Genome Fraction > 80%). We next focused on analysis of animal viruses since the samples were obtained from humans. After removing potential contaminations (see Materials and methods), a total of 28 animal viruses were kept for further analysis. The abundance and positive rate of 28 viruses in samples of three human diseases including Crohn’s disease, Prediabetes, and Ulcerative colitis and healthy people were analyzed. As shown in Figure 5C, a total of 11 viruses were observed in the healthy people, with the Mongoose picobirnavirus and Human picobirnavirus having the highest positive rate. The virome composition in disease samples was different from that in healthy people. For example, only 9 of 16 viruses in the Crohn’s disease were also observed in the healthy people. For each disease group, there were one or more disease-specific viruses. For example, the Rotavirus A which has been previously implicated in the development of intestinal diseases such as the diarrhea, had high abundances in patients of the Crohn’s disease with a median of 4837 FPKM. The virome composition between Crohn’s disease and Ulcerative colitis was similar, with nine virus species coexisting in both diseases, while the Prediabetes had different virome composition to the Crohn’s disease and Ulcerative colitis.

**Figure 5.**
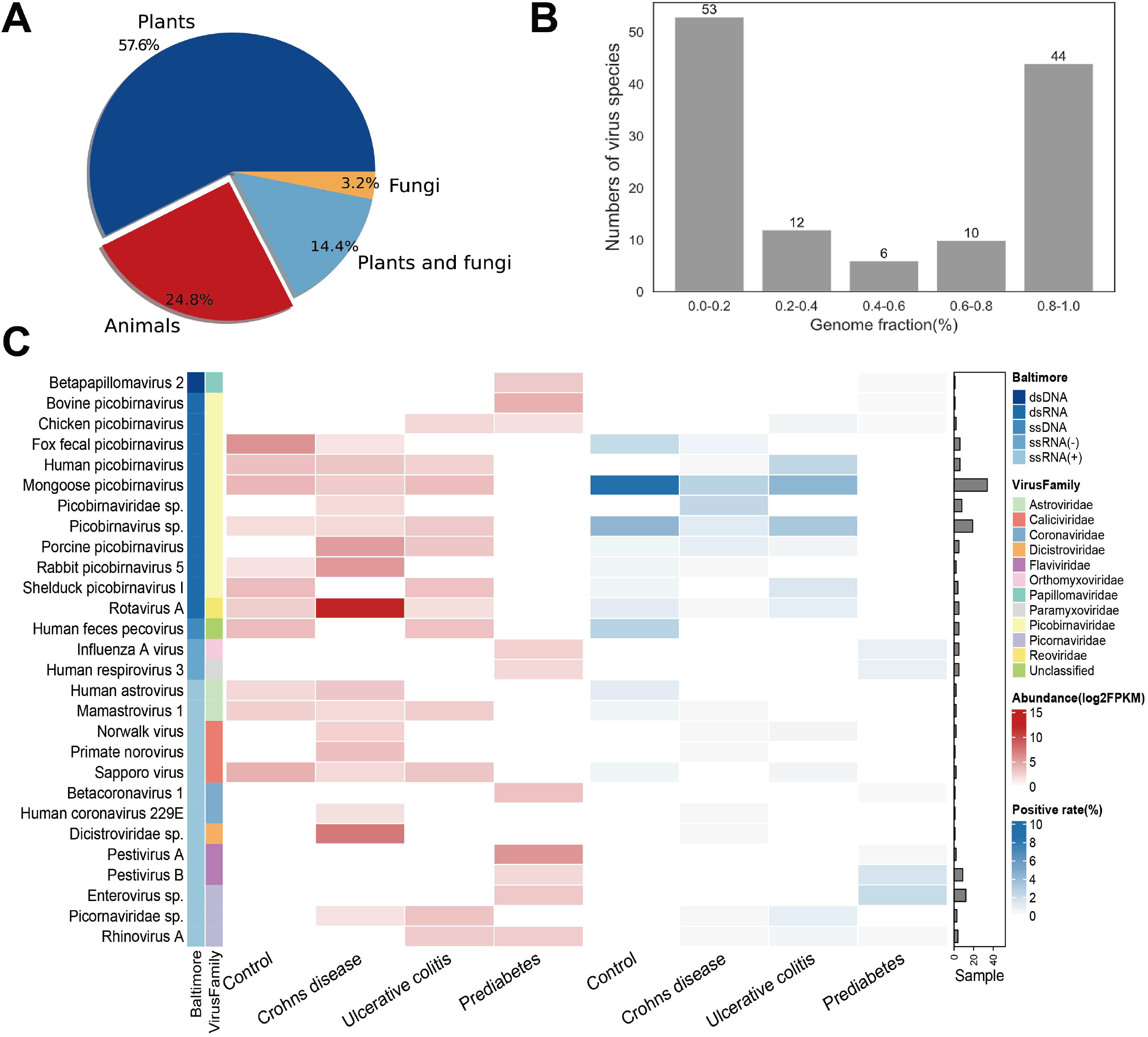
Identification and characterization of viromes in metatranscriptomic data from the HMP dataset. (A) Distribution of assembled viruses based on host type. (B) Number of viruses with a given level of Genome Fraction were assembled. For viruses which were identified in multiple samples, only the most complete genome was used in the analysis. The Genome Fraction was evenly divided into five levels. (C) Abundance and positivity rate of animal viruses in three diseases including Crohn’s disease, Ulcerative colitis and Prediabetes and in the Control group (healthy people). Viruses were organized by groups in the Baltimore classification system and by the viral family. The virus abundance and the positive rate of viruses were shown in the left and central heatmaps, respectively, according to the figure legend in the topright. The bar plot on the right indicates the number of samples in which the virus was identified.

## Discussion

Numerous methods have been developed to identify virus or assemble virus genome from the NGS data. However, there is still a lack of an effective tool for integrating both the functions of virus identification and genome assembly. This study presented an one-stop tool named VIGA for virus identification and genome assembly from the NGS data. It has been shown to assemble more complete virus genomes than its competitors in both the metatranscriptomic and metagenomic data, and also perform well in assembling virus genomes at the strain level. It has also be used to characterize the virome in metatranscriptomic data from the Human Microbiome Project. Overall, it should be an effective tool for virus identification and genome assembly in virome studies.

Compared to previous assemblers, VIGA assembled more complete virus genomes in both metatranscriptomic and metagenomic data. This may be because VIGA integrated both reference-based and *de novo* assembly tools. The *de novo* tools such as Trinity and Haploflow can be used to assemble genomes without dependence on reference genomes. Compared to *de novo* tools, VIGA integrated the reference-based tools which have been reported to assemble more accurate and complete genomes than *de novo* ones^[39]^. Compared to reference-based tools such as MetaCompass, VIGA could automatically select reference genomes from the library of virus reference genomes. Besides, RagTag was used to correct assembly errors for the genomes assembled by MetaCompass. Taken together, VIGA integrated both the advantages of reference-based and *de novo* assembly tools.

Both Haploflow and VirGenA have been designed for strain-resolved assembly of viral genomes. Unexpectedly, VIGA can also be used to assemble virus genomes at the strain level with excellent performances. Compared to Haploflow, VIGA assembled viral genomes with similar precision, higher completeness yet a little lower accuracy on the HIV and HBV datasets. While compared to VirGenA, VIGA assembled more complete viral genomes with higher accuracy. The ability of VIGA on separating mixtures of virus strains may be due to the comprehensive library of virus reference genomes as genome sequences of each virus species were clustered at 100% level. However, for novel viruses without reference genomes in the virus reference genome library, it may be difficult to separate and assemble virus genomes at the strain level.

There were some limitations to the study. Firstly, VIGA can only assemble genomes for known viruses. For novel viruses without reference genomes, it is difficult to assemble accurate genomes by VIGA. Secondly, the accuracy of VIGA in assembling genomes needs further improvements, especially in regions with low abundance or no transcription such as the 3’ and 5’ ends. Thirdly, VIGA can only be used for eukaryotic viruses which have been studied much and for which a large number of reference genomes are available. Fourthly, VIGA can only be used on paired-end NGS data as MetaCompass only accepts such data as input.

## Conclusion

Overall, this study developed an one-stop tool called VIGA for eukaryotic virus identification and genome assembly from the NGS data. It can be used in assembling virus genomes from both metatranscriptomic and metagenomic data and be used in separating virus strains from mixtures. It would help much in identification and characterization of viromes in future studies.

### Key Points

- VIGA is an one-stop tool for eukaryotic virus identification and genome assembly from next-generation sequencing data.
- VIGA outperformed its competitors in assembling more complete virus genomes, even at the strain level, when evaluated on multiple simulated and real virome datasets.
- VIGA can be used to analyze the metatranscriptomic and metagenomic data.
- VIGA is freely available for public use on the GitHub platform at https://github.com/viralInformatics/VIGA.

## Availability of data and materials

All data used in the study are available in public databases. The codes for VIGA is public available at https://github.com/viralInformatics/VIGA.

## Funding

This work was supported by the National Key Plan for Scientific Research and Development of China (2022YFC2303802) and National Natural Science Foundation of China (32170651).

## Acknowledgements

We thank members in PengLab for helpful discussions on the manuscript.

## Competing interests

The authors declare that they have no competing interests.

## Biographical note

Ping Fu and Yifan Wu are PhD students in College of Biology, Hunan University

Zhiyuan Zhang is a master student in College of Biology, Hunan University

Ye Qiu and Yirong Wang are associate Professors in College of Biology, Hunan University

Yousong Peng is Professor in College of Biology, Hunan University

## Notes

### Competing Interest Statement

The authors have declared no competing interest.

