## Supplement Table S2_S3 for "VIGA: an one-stop tool for eukaryotic Virus Identification and Genome Assembly from next-generation-sequencing data"

**Table S2. Performance of different assembly tools on the HIV dataset.** #Normalized with the method of Min-Max Normalization.

| Method | Strain Precision (%) | Normalized Strain Precision^#^ | Genome Fraction (%) | Normalized Genome Fraction^#^ | Mismatches per 100 kbp (bp) | Normalized Mismatches per 100 kbp^#^ |
| --- | --- | --- | --- | --- | --- | --- |
| VIGA | 100 | 1.00 | 98.20 | 1.00 | 2787.00 | 0.93 |
| MetaCompass | 50 | 0.18 | 60.45 | 0.00 | 1463.00 | 0.96 |
| VirGena | 100 | 1.00 | 76.14 | 0.42 | 2766.80 | 0.93 |
| Trinity | 39 | 0.00 | 97.19 | 0.97 | 39585.60 | 0.00 |
| Haploflow | 100 | 1.00 | 93.36 | 0.87 | 33.70 | 1.00 |

**Table S3. Performance of different assembly tools on the HBV dataset.** #Normalized with the method of Min-Max Normalization.

| Method | Strain Precision (%) | Normalized Strain Precision^#^ | Genome Fraction (%) | Normalized Genome Fraction^#^ | Mismatches per 100 kbp (bp) | Normalized Mismatches per 100 kbp^#^ |
| --- | --- | --- | --- | --- | --- | --- |
| VIGA | 100 | 1.00 | 99.91 | 1.00 | 1890.65 | 1.00 |
| MetaCompass | 100 | 1.00 | 99.91 | 1.00 | 2397.20 | 0.64 |
| VirGena | 100 | 1.00 | 46.12 | 0.00 | 1942.18 | 0.96 |
| Trinity | 74 | 0.00 | 73.48 | 0.51 | 3306.42 | 0.00 |
| Haploflow | 100 | 1.00 | 91.19 | 0.84 | 2355.00 | 0.67 |
